# A generalisable framework to inject distance information into Alphafold-like structure predictors

**DOI:** 10.64898/2026.07.02.736010

**Authors:** Claudio Mirabello, Björn Wallner, Vladislav Orekhov, Björn Nystedt, Nicholas Pearce

## Abstract

Structure prediction methods are now highly successful at predicting three-dimensional structures from sequence. However, it is still often desirable to supplement these methods with additional external priors on pairwise distances in the structures. We present a general method for injecting prior information into AlphaFold-like structure predictors by biasing the pair representation to produce desirable features in the distogram, which are then reflected in the structures. We demonstrate this approach to: sample alternate states by selectively pushing or pulling mobile amino acid pairs; integrate NMR NOESY data with structure prediction; and improve the success of protein-protein and protein-ligand complex prediction. We demonstrate that this approach is applicable both to AlphaFold 2 and a reproduction of AlphaFold 3 (OpenFold3).

*resTrain* is open source, available to all users on GitHub and as a Colab notebook: github.com/clami66/resTrain

## 1 Introduction

It has now been over five years since the publication of AlphaFold 2[1] (AF2), which marked a step-change in our ability to predict three-dimensional protein structures from sequence. In this new era of structure prediction, we continue to see the development of deep-learning structure prediction approaches, with moves to diffusion-model-based approaches by the biggest players in the field and the incorporation of RNA, DNA and small molecules into their predictions, notably with AlphaFold 3 (AF3) and RoseTTAFold All-Atom [2, 3].

Alongside these core methodological developments, scientists have explored how to perturb existing methods to produce output structures with desirable features. One such feature is the generation of protein structural ensembles that explore multiple conformations of proteins that involve the reorientation or reorganisation of large parts of the molecule [4, 5]. A common strategy for exploring this conformational heterogeneity is to modify the input multiple sequence alignment (MSA), whose patterns of evolutionary conservation are translated into interactions between different residues in the protein. By clustering, sampling, or masking the MSA input, some evolutionary patterns are enhanced or degraded, changing the predicted interactions, leading to globally different structures [4, 5]. These approaches are largely undirected perturbations of the prediction process, typically requiring the generation of large numbers of models in the hope that at least one sample will present the desired features. A number of methods combine this sampling-and-select approach by scoring models against experimental information, for example to recover multiple protein states by scoring against Nuclear Magnetic Resonance (NMR)-derived data with AlphaFold [6] and AlphaFlow [7].

Other, more focused approaches use *inductive learning* by re-training or fine-tuning AlphaFold to accept extra inputs that encode restraints of various kinds; most of these methods are based on open implementations of AF2 [8, 9], such as AlphaLink [10, 11] and DEERFold [12], though AlphaFold 3 reproductions have been released that can incorporate experimental information in the form of distance restraints. Chai-1 [13] has been trained to accept extra input features in the form of maximum distance restraints between token pairs, or to label single residues as epitope residues (pocket conditioning). Protenix [14] also allows adding restraints of this kind, although the underlying mechanism has not been described in its technical report [14]. These pretrained methods are convenient for users, as they will not need further training or finetuning, but they usually treat the experimental inputs as “soft restraints”, meaning that there is no guarantee that they will actually influence the outcome.

Conversely, other methods fit predictions to restraints by *transductive learning* patterns by optimising network parameters separately for each target. As the optimisation loss decreases, these transductive approaches produce predictions that better satisfy the restraints. Notable examples here are ROCKET [15] and Distance-AF [16].

Here, the restraints are more likely to be incorporated in the predictions whenever the loss term converges, but current solutions suffer from drawbacks that may limit their usability. In ROCKET, for example, memory constraints limit the size of the inputs to single-chain proteins with a maximum of around 500 residues on currently available GPUs. Similar limitations are also found on Distance-AF (tested on a maximum of 1500 residues, single-chain only). Furthermore, the additional neural network layer trained in ROCKET is used to manipulate internal representations within the MSA space, so the method is less likely to be effective for proteins with shallow MSAs, or possibly with multimers. In Distance-AF, which is the approach that is most similar to what we propose in this work, several amino acid pair distance restraints are needed to obtain the desired result.

Few of the above methods are applicable to both monomers and multimers, with some being limited to the former (ROCKET, Distance-AF, AlphaLink) and some to the latter (Chai-1, Protenix). AlphaLink2 has expanded AlphaLink to work with multimers, but only at fixed distance thresholds (25 Å between C_*α*_ atom pairs).

## 2 Results

In this work, we describe *resTrain*, a new flexible framework for integrating experimental information in the form of distance probability distributions between pairs of amino acids in proteins and protein complexes. It is a transductive approach consisting in the optimisation of a bias term to the pair representation in AF2 or AF3-based predictors. The bias term is optimised through a gradient descent procedure to minimise the difference between the desired distance probability distributions and the predicted distogram. We show that optimising distograms alone indirectly enforces the restraints in the structures, so there is no need to apply extra loss terms to the structural module outputs. Using probability distributions instead of fixed targets allows the shape of the target distribution to encode the degree of confidence on any given distance, and makes it possible to represent uncertain distances between atoms, even where those atoms are not explicitly modelled by AlphaFold, such as for H atoms in NMR data.

We implement *resTrain* on AF2 and AF2-Multimer (*AF2-resTrain*) and, as a proxy for AF3 due to licensing restrictions, on OpenFold3 [17] (*OF3-resTrain*). We test *resTrain* in a number of applications, showing that it performs favourably when compared to other state-of-the-art tools. Using different distogram targets, *resTrain* can: alter pair distances to produce multiple conformations for multi-state proteins; integrate Nuclear Overhauser Effect (NOE) data from NMR experiments to guide structure predictions; and integrate sparse or approximate information about interfacial residues to improve modelling of protein-protein and protein-ligand complexes. We also show that it overcomes several of the limitations found in other integrative tools, as a single pair distance is often sufficient to produce the desired results in a handful of gradient descent steps (needing mere minutes of GPU time). Furthermore, resTrain works both on monomers and multimers, as restraints can be enforced both within and across protein chains, and cropping of inputs and outputs during gradient descent permits the optimisation of larger proteins. While we have limited our applications to two prediction methods (AF2 and OF3), it can be easily implemented on any distogram-based prediction pipeline and is thus a generally applicable approach to guiding structure prediction methods.

*resTrain* is open source, available to all users on GitHub and as a Colab notebook (https://github.com/clami66/resTrain).

### 2.1 Driving predictions through distogram restraints

Distograms were introduced in the first version of AlphaFold [18]. A distogram is a set of probability distributions for the distances between *C*_*β*_ atoms for all pairs of amino acids in a structure. In AlphaFold, these distributions of *C*_*β*_ *− C*_*β*_ distances are classified into 64 discrete distance bins, ranging from [2.31, 2.62) Å in the first bin to [22, ∞ ) Å in the last bin. In AF2 and AF3, distograms are auxiliary outputs obtained from the pair representation output of the EvoFormer (PairFormer in AF3) trunk (Fig. 1) and are not directly involved in the structure prediction step. However, distograms and structures are both influenced by the same pair representation. Thus, we hypothesised that manipulating the pair representation in the EvoFormer to increase the distogram probability for lower distance bins would also realise smaller distances between pairs of residues in the final structure. If that were the case, manipulating distograms would be a simple and elegant way of integrating distance restraints in the prediction of structures.

**Fig. 1:**
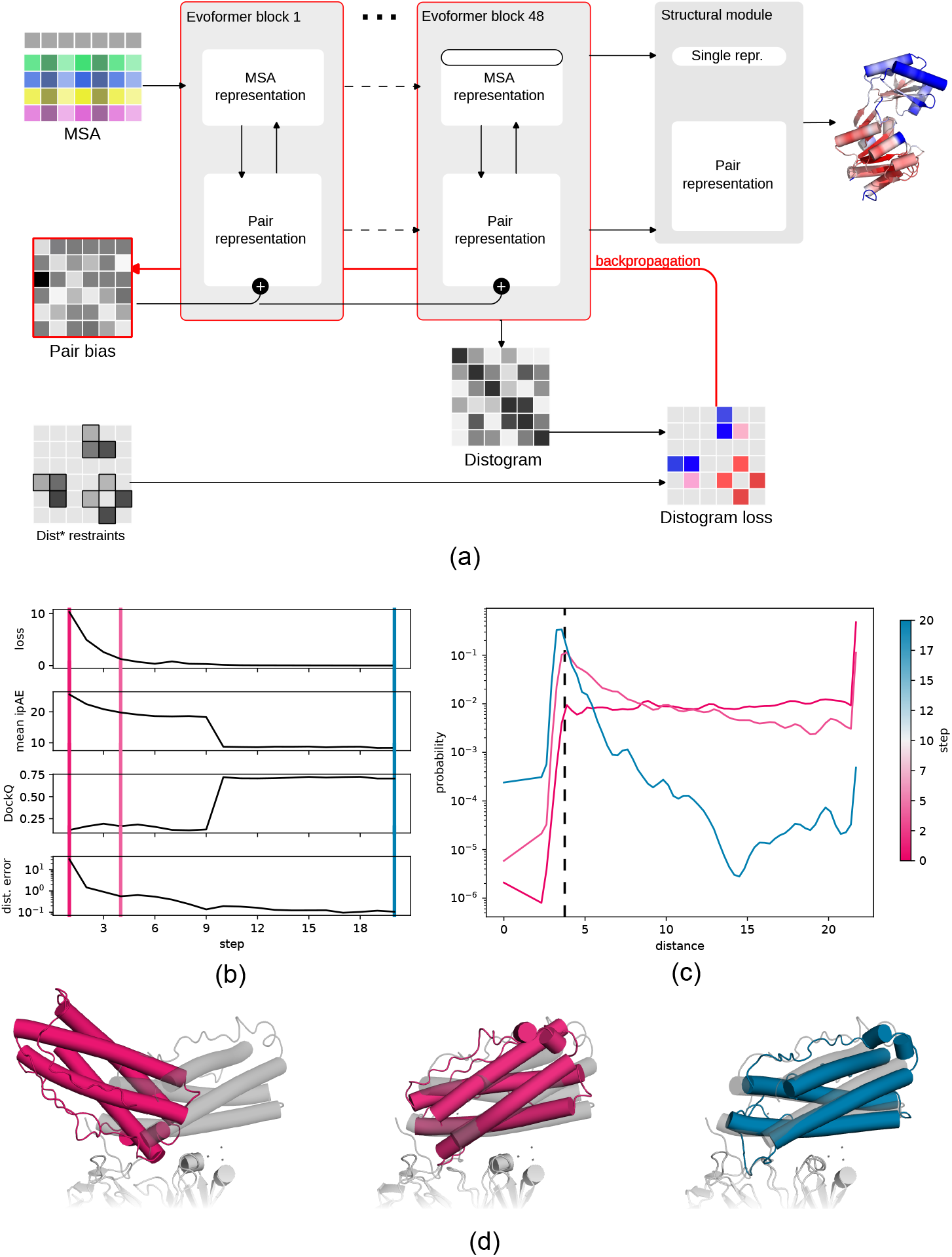
(a) Flowchart of the *resTrain* method. The distogram loss is backpropagated through the Evoformer modules to a common pair bias, which is updated during training. (b-d) Antibody-antigen target example (PDB 7N0A). The structure is predicted in the wrong conformation by AlphaFold2, but is predicted correctly by *resTrain* when a single distance restraint is applied between the antigen (GLY124) and CDR H3 (HIS31) at 3.74 Å distance. (b) Various quantities during *resTrain* optimisation: distogram restraint loss, mean ipAE for predicted structure, DockQ interface score, and actual distance error of the restrained interaction. Step 0 corresponds to the original AF2 prediction. After 10 optimisation steps, updating the restraint pair bias (RPB), the predicted structure switches to the correct conformation, as reflected by a step change in the DockQ score. Marked positions 1, 4 and 20 correspond to elements in (c) and (d). (c) Evolution of the distogram for GLY124-HIS31 after training steps 1, 4 and 20. What begins as a broad distribution across bins becomes concentrated around the target value, 4Å. (d) Evolution of the predicted structures after training steps 1, 4 and 20. The deposited structure is shown in semi-transparent grey, and structure colours correspond to line colours in (b) and (c).

In our implementation (see methods for full details), the user inputs a set of distance restraints for defined amino acid pairs. These restraints can either be probability distributions (using the binned distances described above), or maximum distances. Using appropriate loss functions for the type of distribution, a gradient descent procedure iteratively scores the predicted distogram against the restraints and back-propagates the resulting loss through the network. Gradients are not applied to the neural network itself, but to a new auxiliary input that we call the *Restraint Pair Bias* (RPB). The RPB is an extra bias term to the pair representation in every Evo-former block. Thus, the RPB is optimized so that the predicted distogram matches the restraints.

An illustrative example is shown in Fig. 1b-d. In this example, a single correct distance restraint at the interface between an antibody-antigen complex is enough to drive a correct prediction in 10-20 optimisation steps (PDB ID: 7N0A; 3.74Å restraint between antigen GLY124 and CDRH3 HIS31). As the loss decreases, there are improvements in both objective quality measures (DockQ score, distance error of restraint) and predicted quality measures (mean interfacial predicted Aligned Error, ipAE) (Fig 1b). Meanwhile, distogram probabilities for the restrained pair progressively increase around the selected distance (Fig 1c). Finally, the changes in the distogram are reflected in the structural output itself (Fig. 1d). The distogram changes are not limited to the defined pair, but propagate to the whole interface (Fig. S1).

In the following sections, we demonstrate the injection of both exact and approximate distance restraints in various application areas: prediction of alternate conformational states; NMR structure generation from NOE data; the prediction of hard multimeric complexes; in-silico epitope scanning; and finally ligand docking.

### 2.2 Alternate conformational states and trajectories

Many proteins are dynamic and can change conformation in order to perform their function. Channel proteins, for example, can adopt multiple states as they regulate access to a cell through its membrane. Whenever a protein structure has multiple states, AlphaFold often shows a preference for one of those states and seldom (in some cases never) predicts the other states [4].

Here, we show that, if information about labile interactions in a structure is known, it can be leveraged in the *resTrain* framework to direct the generation of ensembles of conformers. For protein that are always predicted in a “closed” conformation, pushing apart the right pair of residues or breaking the right contacts can force the sampling of other conformations. From the Cfold dataset [19], we selected the proteins where two conformations have been deposited in the PDB. For each structure, we select the most mobile pair of residues for each and run a series of *AF2-resTrain* experiments enforcing inter-pair distances from 3-22+Å (see Methods). Pulling or pushing the right pair of residues proves to be as good or better than random sampling in AlphaFold to retrieve the other state and, as a directed approach, can produce good results with tens, rather than thousands, of predictions (Fig. 2a,b). For several proteins in the Cfold set, we also see that sweeping the distance between the selected pair of residues can generate hypotheses about the trajectory between the known states, as well as proposing previously unseen states (Fig. 2c).

**Fig. 2:**
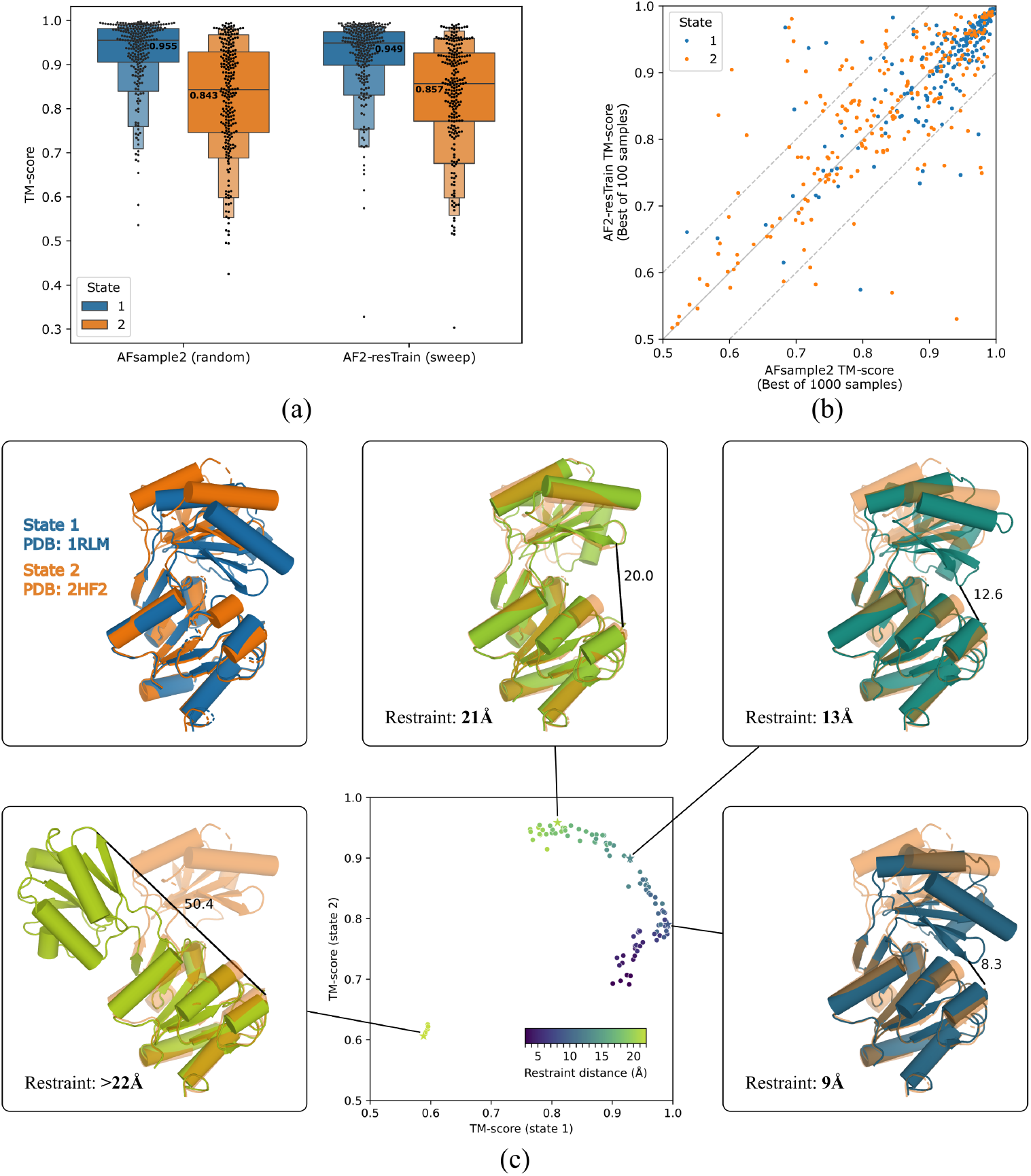
Applying springs to pairs of amino acids drives conformations between alternate states. (a,b) Comparison of AFsample2 (“random”) and resTrain (“restrained”) for the generation of multiple states for 238 reference pairs from the Cfold dataset. For each target, 1000 structures are generated with AFsample2, while 100 structures are generated for *resTrain* (20 restraint distances from 3-22Å for each of the five AF2 models). (a) Distribution of best TM scores to each state across the example targets. (b) Same data as (a). *resTrain* produces significantly better TM scores (Δ*TMscore >* 0.1) for 31 states, compared to 16 for AFsample2. (c) Predicted structures from *resTrain* sweep for PBDID 1RLM (state 1, blue) and 2HF2 (state 2, orange), plotted by their TM-score to each state. *resTrain* predictions are coloured by restraint distance; the observed distance is shown for each structure. Highlighted structures are the deposited structures (top-left), the highest TM score to state 2 (top-centre), an intermediate state between state 1 and state 2 (top-right), the highest TM score to state 1 (bottom-right) and a structure with the lowest TM scores to both observed states (bottom-left). State 2 is shown semi-transparent in all images for reference. Predicted structures form a trajectory between the two known states, as well as proposing new previously unobserved structures.

### 2.3 Integration of predictions with NMR restraints

Structures determined by nuclear magnetic resonance (NMR) spectroscopy are not directly imaged, in contrast to macromolecular crystallography or cryo-electron microscopy. Instead, NMR NOESY experiments measure atomic-level interactions which are converted into structural restraints (e.g. interatomic distances). In structure calculation with CYANA [20], a popular NMR software, restraints are combined with torsion-angle dynamics with simulated annealing[21, 22] to generate an ensemble of structures.

To see if NOESY data could be used to guide structure predictions through the distogram, we applied *AF2-resTrain* to the ARTINA set [23], for structures where NOESY data was available. The NOESY data provides a maximum distance for a pairs of hydrogen atoms. Because the distogram predicts *C*_*β*_-*C*_*β*_ distance distributions, we constructed a database of H-H distances and corresponding *C*_*β*_-*C*_*β*_ distances from the PDB [24, 25]; this data is used to derive *C*_*β*_-*C*_*β*_ distance distributions from given H-H distances (see methods), which are then used as the target for resTrain. We compare AF2, AF2-resTrain, and the deposited PDB structures; for each, we choose as a representative the structure with the lowest aggregated violations (the sum of restraint violations across the structure).

Application to 90 structures from the ARTINA set shows that optimising the distogram with *AF2-resTrain* leads to decreased violations for 88% of structures, reducing the aggregated violations by 25%, and decreasing the number of violated restraints by 19%, compared to AF2 predictions (Fig. 3a-b, Fig. S2). The most significant changes are for target 2L82, where AF2 swaps two of the core beta sheets; *AF2-resTrain* corrects this error (Fig 3a,b,e-f).

**Fig. 3:**
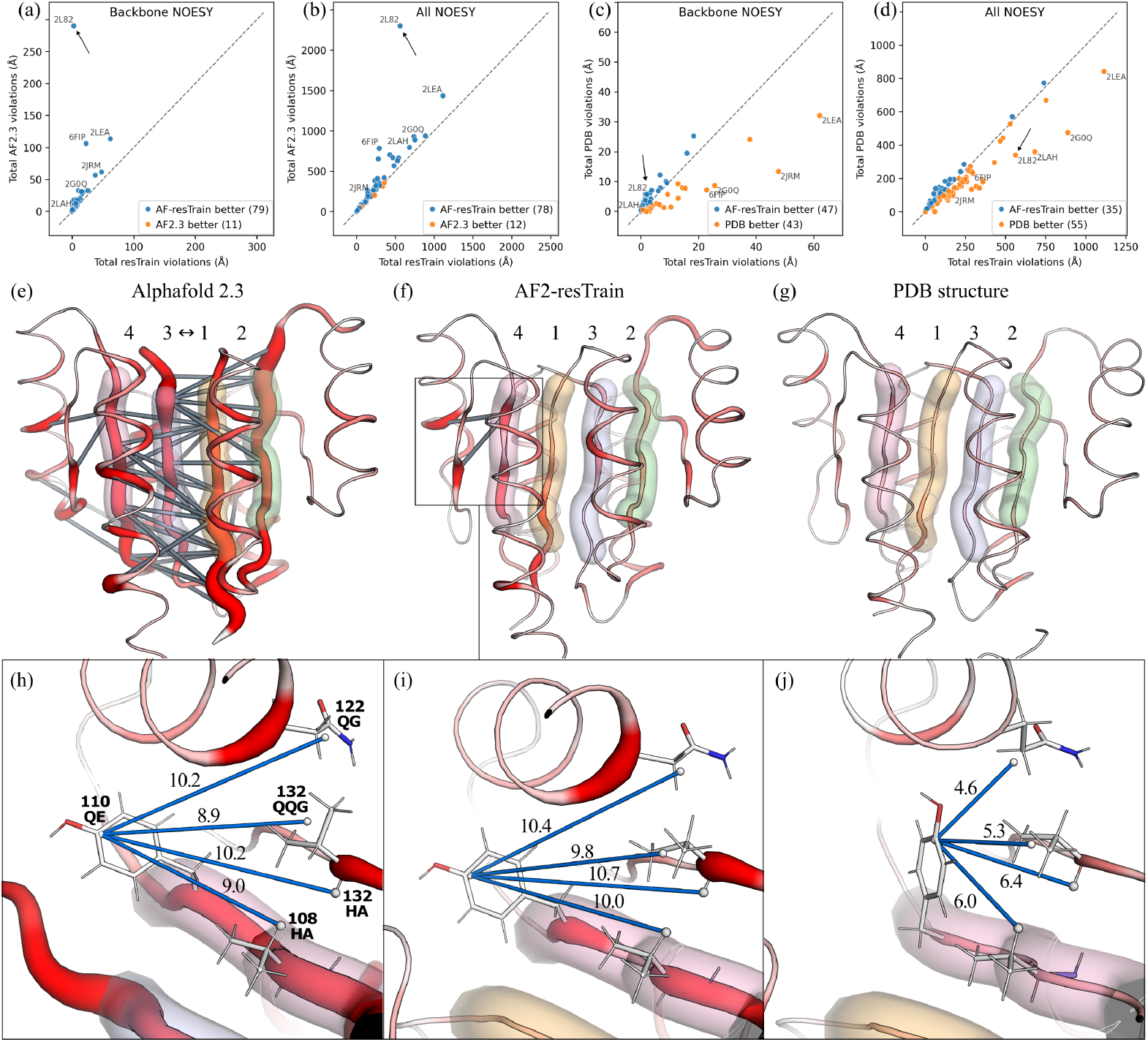
Effect of NOE restraints on prediction of structures in the ARTINA dataset. Total NOE violations for (a) backbone-backbone NOEs and (b) all NOEs for AF2-*resTrain* predictions compared to AF2 predictions. *resTrain* models demonstrate significantly fewer violations, showing that the optimisation of the distogram has had the desired effect on the predicted structure. Total NOE violations for (c) backbone-backbone NOEs and (d) all NOEs for *AF2-resTrain* predictions compared to deposited PDB structures. The performances of the two approaches are comparable, though the deposited NMR structures are clearly better at satisfying sidechain interactions. (e-j) Example 2L82, which is improved markedly by use of NOE data. Cartoon representation of the lowest-violation model for (e) AF2 (f) AF2-resTrain and (g) deposited PDB ensemble. Putty width and colour reflect the largest NOE violation involving that residue. Violations larger than 5Å are shown as grey bars connecting *C*_*β*_ atoms of the two residues. (e) The AF2 prediction contains a large number of violations due to an incorrect arrangement of two core Beta-sheets (coloured semi-transparent tubes; numbered 1-4 by sequence position). (f) After application of resTrain, the position of the two central strands is swapped. (g) The deposited PDB structure also shows the correct arrangement and very few violations. “Backbone” interactions are those that involve H, HA, HB or QB atoms. (h-j) Zoom in of largest remaining violation in AF2-resTrain structure. Atoms involved in NOEs are labelled and shown as spheres. NOE restraints are shown as blue bars, but annotated with the observed distance. AF2 (i) resTrain and (j) PDB structure. The PDB structure satisfies the NOE restraints with a TYR conformation that shifts the helix containing residue 122 upwards.

*AF2-resTrain* structures are broadly on par with the deposited PDB structures (Fig. 3c-d): PDB structures have 17% fewer aggregated violations, while *AF2-resTrain* structures have 6% more satisfied restraints (Fig. S2). *AF2-resTrain* performs worse on those structures where the deposited structure also scores poorly (violation totals over 500, Fig 3d); it is not clear whether the models (or potentially, the derived restraints) are of high enough quality to derive meaningful differences between them, and a reassessment of the underlying assignments is outside the scope of this project. From inspecting the largest remaining violations, it is clear the largest errors in *AF2-resTrain* models are caused by incorrect sidechain conformations, which are only indirectly affected by optimising the distogram (e.g. Fig 3h-i).

Our results here show that distogram optimisation may be an efficient way of steering future “sample and select” approaches for AF2-like structure predictors, where experimental restraints are available.

### 2.4 Improving predictions of hard multimeric complexes

Using the *resTrain* approach, we can enforce approximate restraints, such as a maximum distances between residues; in the loss function these restraints rewards probability mass under the restraint threshold, while being agnostic to its exact location. This feature is particularly appropriate for the prediction of protein complexes, where interfaces may be known, but not necessarily specific interactions. Additionally, since they are generally larger in size, protein multimers could exceed the amount of GPU RAM available during the gradient step. This is resolved in *resTrain* training by cropping the inputs around restraints provided by the users.

We demonstrate this mode of *resTrain* using 25 recently deposited structures of antibody-antigen complexes. These complexes are hard to predict as the interfacial residues do not co-evolve as with native protein-protein interactions, meaning that interactions are not strongly encoded in the MSA. For a given complex, we enforce a single, approximate distance restraint capped at 8Å between the antigen epitope and the heavy antibody chain. We find that it is not necessary to restrain the light chain, as the interface between the two antibody chains is always correctly predicted regardless of the location of the heavy chain.

This single approximate restraint drives the success of complex prediction from 44% for AF2 to 72% for *AF2-resTrain* (11 and 18 correct predictions, respectively; Fig. 4a,b). This is a striking result, as it shows providing a single imprecise contact between chains can be enough to fully determine the geometry of the interaction. Furthermore, the correlation between AF2 predicted interfacial quality (*ipTM* ) and the DockQ score is generally good for the same targets (Fig. S4,S5). Moreover, predictions with the lowest distogram loss are not necessarily those with best predicted quality (Fig. S4; e.g. targets 7SO7, 7ZXK, 8DKE). This is evidence that the *resTrain* approach does not bias the predicted quality to prefer models that better fit the restraints, but rather, that the predicted quality measures are still reliable at picking the best models.

**Fig. 4:**
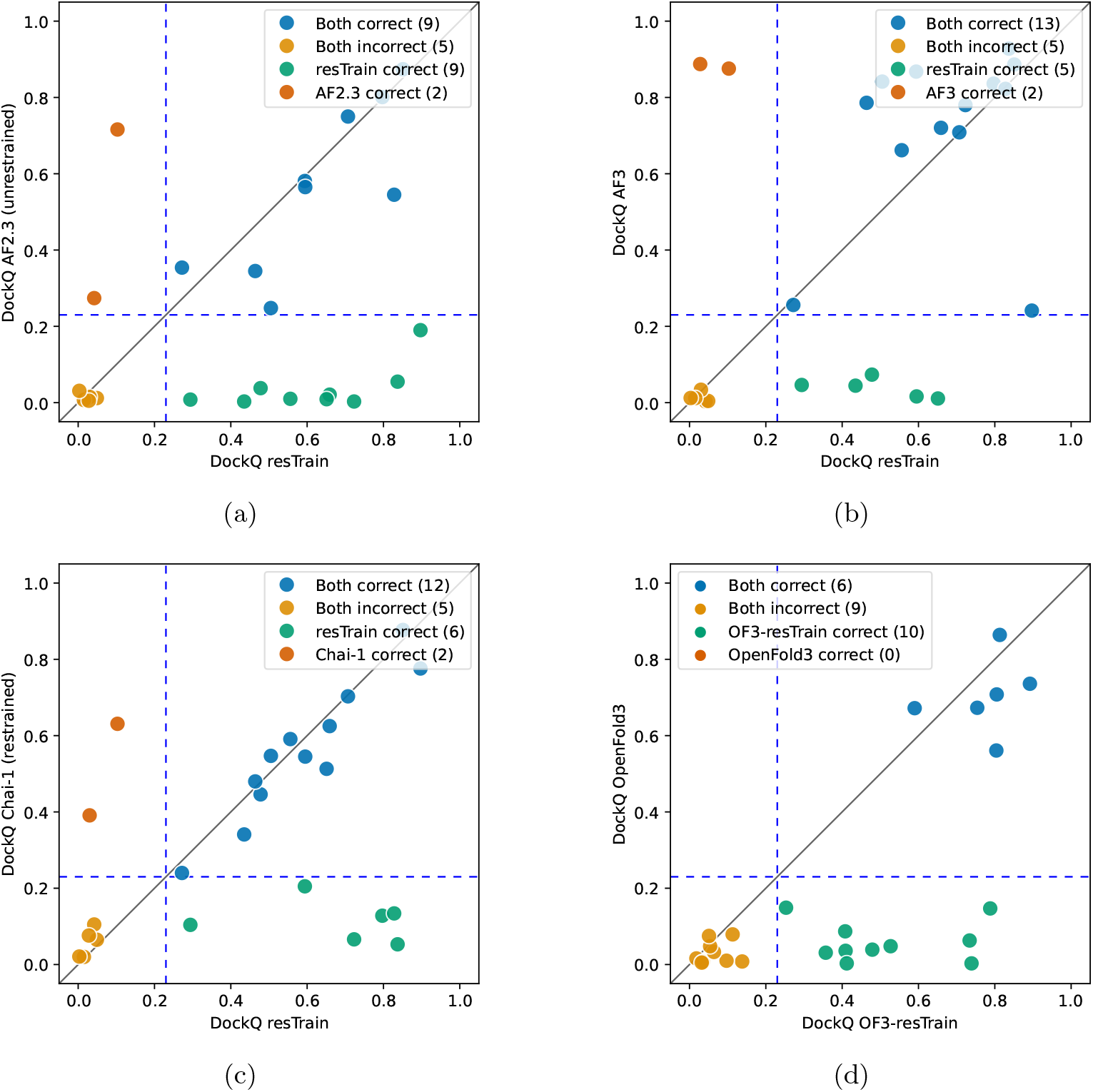
We compare results obtained by running (a) AlphaFold 3, (b) Unrestrained AlphaFold-Multimer (v2.3) and (c) restrained Chai-1 against AF2-resTrain on a set of 25 “hard” protein multimers. If only one pair of interfacial residues is known (approximate distance within 8Å), the gradient descent procedure can correctly (antibody-antigen DockQ score above 0.23) capture the resulting interaction for 18 out of 25 target with relatively little sampling (20 seeds, 5 predictions per seed). This compares favorably to running more extensive sampling with AlphaFold 3 (15 out of 25 targets correctly predicted with 1000 seeds, 5 predictions per seed), AlphaFold-Multimer (11 out of 25) and Chai-1 when using the same set of restraints with similar levels of sampling (15 out of 25). (d) Implementing resTrain on AF3-based architectures (OpenFold3 in this case) similarly improves results.

Moreover, the *AF2-resTrain* predictions compare favourably even against AF3, which is currently the best tool at predicting antigen-antibody complexes [17], even as sampling is much wider with AF3 (1000 seeds, 5 predictions per seed as suggested by the authors) compared to *AF2-resTrain* (20 seeds, 5 predictions per seed).

In order to compare against a state-of-the-art method that allows external restraints, we also compare to Chai-1, which is based on the same architecture of AF3 but can accept additional distance restraints (maximum distances between pairs of residues). Using the same set of single-pair restraints and the same number of predictions for Chai-1 as for *AF2-resTrain*, we still observe superior performance for *AF2-resTrain*, with 14 correct predictions for Chai-1 (Fig. 4c).

Lastly, we applied *OF3-resTrain* (a modified version of OpenFold3-preview), to explore whether diffusion-based AlphaFold3-style architectures also benefit from the distogram modifications. This proves to be the case, with 16 correct predictions for *OF3-resTrain* compared to only 6 for OpenFold3; this performance is still notably less than *AF2-resTrain*.

### 2.5 In silico epitope scanning

Generating AlphaFold predictions – even in large numbers – quite often results in few clusters of very similar structures. Using *resTrain*, however, it becomes possible to systematically sample areas that might not be otherwise explored. In the case of protein dimers, for example, we can systematically sample the space of possible interfaces by restraining possible cross-chain pairs of residues to be in proximity. This simplifies with additional knowledge of the binding interfaces, as in the case of antigen-antibody complexes, where the antibody interacts with the antigen through the Complementary-Determining Regions (CDRs).

We demonstrate this “epitope scanning” approach on one of the smaller antigens from the antibody dataset (PDB ID: 7W71). We pick a single residue in CDR H3 (SER101) and run 93 different gradient descent experiments (approximate distance restraint, within 8Å) to each of the 93 antigen residues in turn. As for the previous experiment, we generate 100 predictions (20 seeds, 5 predictions per seed) for each of the 93 experiments, for a total of 9300 structures. The top-ranked structure by ipTM correctly predicts the interface (DockQ score: 0.74, Fig. S6).

While not exhaustive, this test shows the potential of epitope scanning in our framework, and how the predicted quality scores in AlphaFold are still valuable for identifying the most promising hypotheses.

### 2.6 Protein-ligand complexes

Implementing the *resTrain* approach within AF3-like predictors expands applicability to small molecules. Here, distance restraints could be provided to target known binding pockets during virtual screenings. We briefly test whether *OF3-resTrain* can correctly model protein-ligand interactions on a single hard target. We select a target (PDB ID: 8*OV* 7) from the *Runs N’ Poses* benchmark set [26] where AF3 has been shown to fail to dock a small molecule ligand against the receptor protein (reported ligand RMSD: 12.6Å). We apply *OF3-resTrain* with two restraints for two heavy atoms at either extremity of the ligand (C18, N) against two residues in the pocket (PHE87 and ARG166 respectively). For simplicity, we select the residues that are closest to the corresponding heavy atom in the deposited structure. Both restraints are set to an upper-bound distance of 8Å, significantly higher than the actual distances in the deposited structure of 4.8Å and 5.6Å respectively. The top-ranked model by ipTM out of 100 (20 seeds, 5 predictions per seed) is correctly predicted, with a pocket-aligned ligand RMSD of 1.24Å (Fig. S 7).

## 3 Methods

### 3.1 *resTrain* implementation

#### 3.1.1 Architecture and training protocol

In *resTrain* we add one component to the standard neural network model in AF2: the *Restraint Pair Bias* (RPB). The RPB has shape (*r, r*, 128) where *r* is the number of amino acids in the target protein or protein complex. As such, the RPB has the same shape as the “pair representation” in AlphaFold’s EvoFormer and is used to bias it, being applied before every EvoFormer block. While the NN parameters are not shared across the 48 EvoFormer blocks in AF2, the RPB is the same across all blocks (weight sharing). This reduces the number of learnable parameters at training time while still effectively steering the network. We also find that biasing all EvoFormer blocks is more effecting than biasing e.g., only the first or last blocks. The RPB is the only component of the network that is trained during the gradient descent procedure: all other network parameters are frozen.

The algorithm is summarized in Algorithm 1 and proceeds as follows. At training time, the RPB is initialized to zero. The loss is calculated from comparing the distogram output against the set of input restraints through a masked loss function. The masking is necessary to backpropagate errors only for those pairs of amino acids where the user has provided a distance restraint. The loss functions are defined in a following section.

##### Algorithm 1

Training procedure for AF2-resTrain. New components are highlighted in yellow

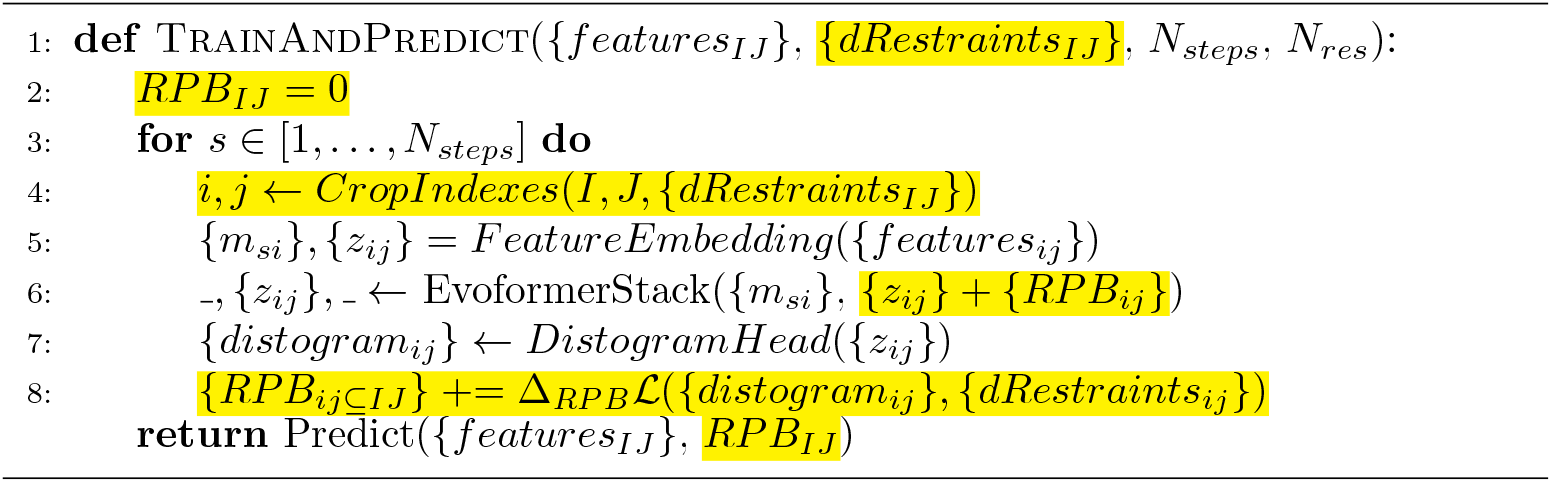

The distogram loss is backpropagated through the network and gradient updates are applied to the RPB. For large proteins, storing the gradients can easily exceed the amount of available GPU RAM, so we implement a modified version of the cropping procedure described in the original implementation of AF2, as follows. A random pair of amino acids is selected from the set of restraints. Then, a square window (384 to 512 amino acids in size in our tests) is defined on the distogram, centered at the selected pair, to extract a subset of rows and columns *i, j* ⊆ *I, J*. The corresponding columns of amino acids are extracted from the MSA representation. The gradients will then flow only to the selected window of amino acid pairs in the RPB. In *AF2-resTrain*, the number of MSA clusters and maximum number of extra MSA sequences have been decreased to 256 each to reduce the need for GPU RAM when training crops of size 512, since the gradient descent procedure becomes more efficient with larger crops. Multiple crops can be extracted and used in a training step in batches, which may speed up convergence in some cases.

One epoch in the gradient descent procedure is made of a fixed number of gradient steps, followed by a single inference step. The RPB learning procedure is quite efficient, often around 5 to 10 gradient steps per epoch (batch size: 1) are sufficient to achieve good agreement between predictions and restraints. In cases where the restraints are many and imperfect, such as in the NMR test described below, performing up to 1000 gradient steps could yield better results. Once the gradient steps are completed, a single, full-size (no cropping) inference step is performed to predict the full structure. For multiple predictions, the RPB is reset to zero and a new RPB optimisation is executed to generate the second prediction, and so on.

In *OF3-resTrain* we adapt the algorithm to the AlphaFold3 architecture. We found that a bias term in the Pairformer only was not enough, and so added a second RPB to the pair representation in the MSA module, in addition to the bias in the PairFormer.

#### 3.1.2 Types of distance restraints and corresponding loss functions

There are several different restraints that may be placed on the distogram, and these each have corresponding loss functions. The restraints we consider in this work are: exact distances, approximate distances (e.g. maximum distances), and distance probability distributions.

Depending on the type of restraint, we therefore use different loss functions:

- Exact distances: Softmax cross-entropy loss (the same loss that was used to train the distogram head in AF2). The appropriate distogram class (distance bin) is set to 1 while all others are set to 0.
- Approximate distances: Sigmoid cross-entropy loss (*AF2-resTrain*). All distogram classes below the upper bound are set to 1 while the other are set to 0. In *OF3-resTrain*, however, we find that the gradient descent procedure is less likely to become stuck in local minima when using a partial-label cross-entropy [27].
- Probability distributions: Kullback–Leibler divergence. Distogram classes are set to the distribution probabilities. Distribution must already be binned appropriately.

### 3.2 Benchmarks and Case Studies

#### 3.2.1 Protein alignments and templates

For all tests, we run MMseqs2 [28] to generate Multiple Sequence Alignments (MSA) following the protocol used in ColabFold [29] and by aligning against Uniref30 (2302) alone. The data pipeline in *AF2-resTrain* has been extended to allow the use of MMseqs2 alignments, while OF3 already uses MMseqs2 alignments from ColabFold’s MSA server. In multimeric tests, MSA pairing has been disabled to disentangle the effect of applying restraints from noisy MSA pairs. For the same reason, templates are disabled in all tests.

#### 3.2.2 Quality measures

We use DockQ [30] to measure the quality of predicted interfaces in the antigen-antibody dataset. The final DockQ score is calculated by averaging individual DockQ scores of the two antigen-antibody interfaces. We exclude the interface between the antibody chains as that is always correctly predicted.

We also use DockQ to calculate the pocket-aligned ligand RMSD in the small molecule ligand docking example.

The quality of monomer conformers is calculated with TM-score [31].

Violations in the NMR analysis are calculated between hydrogen atoms or pseudoatoms as defined in the UPL files. Where multiple atoms are equivalent (e.g. hydrogens in a methyl group), all valid distances are combined using the formula 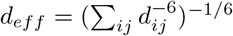

### 3.3 Conformer Set Analysis

For each pair of the 238 pairs of reference conformers in the Cfold dataset, we select a single residue pair to restrain in our experiments. We extract and compare residue pair distances from the two reference structures, and pick the most mobile pair of residues, defined as the pair with highest distance discrepancy across the two reference conformers, provided that the amino acids are in contact (*C*_*β*_ − *C*_*β*_ distance within 8Å) in at least one conformer.

Once a pair is selected, we vary the selected *C*_*β*_ − *C*_*β*_ pair restraint distance between 3 and 22Å(in steps of 1Å). This means that for a single protein, we run 20 different gradient descent experiments in parallel.

We compare our results against those obtained with random sampling by AFsample2 [4] with 20% random MSA masking and 1000 predicted samples (100 predictions for each of the 10 *monomer* and *monomer ptm* models in AlphaFold2).

### 3.4 NOE Restraints

We use the ARTINA benchmark set [23] to test the integration of NOE restraints within our structure prediction framework. This dataset includes 100 NMR structures from the Protein Data Bank. NOE data has been deposited for 94 out of the 100 proteins, and another four proteins were removed as they caused AlphaFold’s relax procedure to crash, making the final dataset 90 proteins. These sets of NOE restraints consist of a number of hydrogen pairs, each associated with a maximum distance. Some interacting atoms are pseudoatoms, formed by averaging the positions of chemically equivalent hydrogens within a residue, but we refer to both hydrogens and pseudoatoms collectively as hydrogens.

To achieve distance probability distributions between pairs of *C*_*β*_ atoms (*C*_*α*_ in the case of Glycine), we constructed a dataset of H-H and corresponding C_*β*_-C_*β*_ distances from the PDB. For a non-redundant set, we use the SCOP class definitions (2022-06-29 release). Hydrogens were then added with REFMAC [32], resulting in 17282 domains. Notably, this data set only contains single domains, as per the typical NMR experiment, and therefore may not generalise to the prediction of inter-domain C_*β*_-C_*β*_ distance distributions; this is beyond the scope of the current application.

Pseudoatoms were added following the CYANA naming convention, with some disambiguations to distinguish between methyl groups and methylene groups (Table S1). Pseudoatoms are placed at the geometric average of their substituent hydrogens. Each hydrogen and pseudoatom are then assigned to a class: each chemically unique pseudoatom forms its own class; chemically equivalent pseudoatoms within residues are grouped together; hydrogens within a pseudoatom take the identity of the pseudoatom plus x (e.g. ALA HB1 maps to MBx); and finally unique hydrogens not part of pseudoatoms are named unambiguously (Table S2).

Catalogued distances are further classified by the sequence separation between the two residues: distance distributions between residues close in sequence space are distinctly different than those from sequentially distant residue pairs (Fig. S3). We observed this signature dissipated after a sequence separation of ±5, leading to larger sequence separations being cached together, i.e., larger sequence separations are mapped to a general distribution.

The observed counts were then binned according to H-H as well as *C*_*β*_-*C*_*β*_ distance. For the *C*_*β*_ bins we use the same binning as that present in the AF2 distogram (64 bins); for Hydrogen we use 48 bins (Table S3). The result of this process is an array of counts corresponding to different sequence separation, H-H, and C_*β*_-C_*β*_ distances for each residue-residue type and hydrogen-hydrogen class (examples: Fig. S3). For a query maximum H-H distance, the appropriate table is identified by residue and hydrogen types, and then truncated at the appropriate H-H limit. If the number of observations in the truncated table is below some minimum, tables from from other residue types are included, starting with the most similar, as measured by the Wasserstein distance, until the minimum number is reached. We then row-wise normalise the distributions to probabilities (such that a probability distribution of *C*_*β*_ distances is obtained for each H-H distance).

The NOE data provides H-H restraints in the forms of maximum distances, which do not, however, imply a uniform probability distribution for interaction distances below this threshold. It is not possible to know this distribution *a priori*, so we derived a prior from observations in the ARTINA set. This data set provides a set of H-H maximum distances, and corresponding observed distances; binning by maximum distance, we obtain a set of distributions of actual distances for each maximum distance.

For the appropriate H-H maximum distance, the prior H-H distribution is used to weight the row-wise normalised counts; these are then summed over the H-H axis, creating a probability distribution of *C*_*β*_ distances for each H-H restraint.

Thus, a C_*β*_-C_*β*_ distance distribution is extracted for each NOE. Where multiple NOEs are present between two residues, these need to be reduced to one C_*β*_-C_*β*_ distribution. Since we do not record information about the correlation between different distances in a residue pair, we apply a coarse naive averaging over the distributions. Functions explored were: multiplicative, arithmetic average, harmonic average and geometric average. The multiplicative reduction approach was observed to produce the best results, though all options are available in the *resTrain* implementation.

### 3.5 Antibody-Antigen Complexes

In this case, we use a subset of recent antibody-antigen complexes gathered from the PDB in [33]. The original dataset includes 71 PDB structures and 166 interfaces in 66 interface clusters. Antibody-antigen complexes are hard to predict correctly as there is no coevolutionary signal across their interfacial residues, and even AF3 correctly predicts around 2 out of 3 such complexes. In order to save computational resources, we selected a subset of structures by keeping only one structure when multiple share the same set of interface cluster keys. In order to test the hypothesis that as little as a single distance restraint is enough to make a good prediction, we also removed complexes with more than one antigen chain. The resulting subset is made of 25 structures.

For each structure, we generated a single distance restraint between the closest pair (lowest *C*_*β*_ *− C*_*β*_ distance in deposited structure) at the interface between the antigen and antibody heavy chain. The restraint was not set to the actual pair distance but rather approximated as a cross-chain contact by capping the distance to 8Å. This is always higher than the actual distance in the deposited structure. This allowed to test a more realistic scenario where the user might only know the approximate location of the antigen epitope.

## Acknowledgements

This work was funded by SciLifeLab as a Technology Development Project (BeyondFold). CM is financially supported by the Knut and Alice Wallenberg (KAW) Foundation as part of NBIS at SciLifeLab. BW is financially supported by KAW as part of the WASP-DDLS joint program. NP is supported by the SciLifeLab & Wallenberg Data Driven Life Science Program (grant: KAW 2020.0239). The computations were enabled by the Berzelius resource provided by the Knut and Alice Wallenberg Foundation at the Swedish National Supercomputer Centre. The authors would like to thank Erik Ylipää and Dr. Daniel Malmodin for helpful discussions and input during the development of this study.

