## Supplementary information for "A generalisable framework to inject distance information into Alphafold-like structure predictors"

July 2, 2026

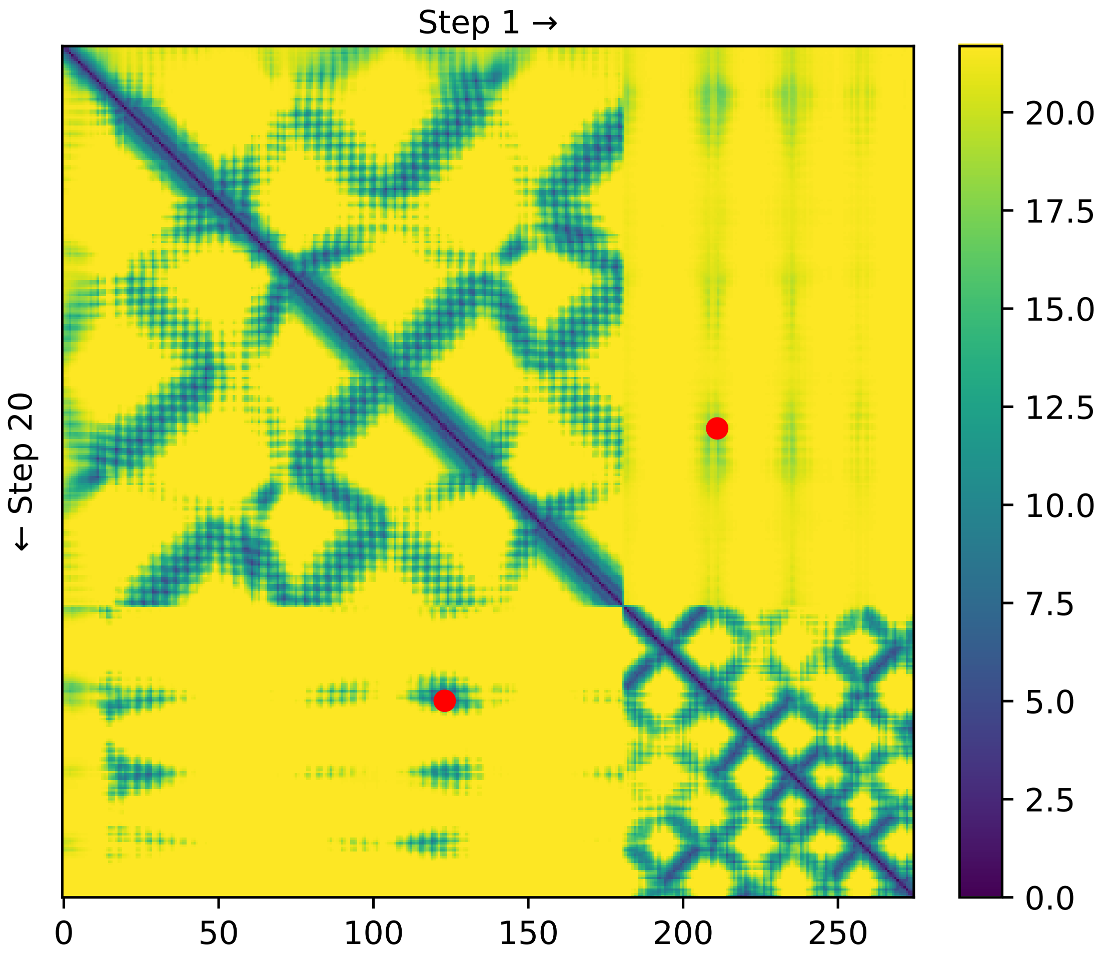


Figure 1: Example of distogram evolution during the gradient descent for dimeric structure shown in main text Fig. 1 (b-d). As the gradient descent procedure optimises the loss on a single restraint at the dimeric interface, the distogram prediction changes from its initial state (step 1, upper triangle) to its final state (step 20, lower triangle). The distogram changes not only for the single contact provided as restraint (red dot), as the learned RPB causes changes to propagate to the whole interface.

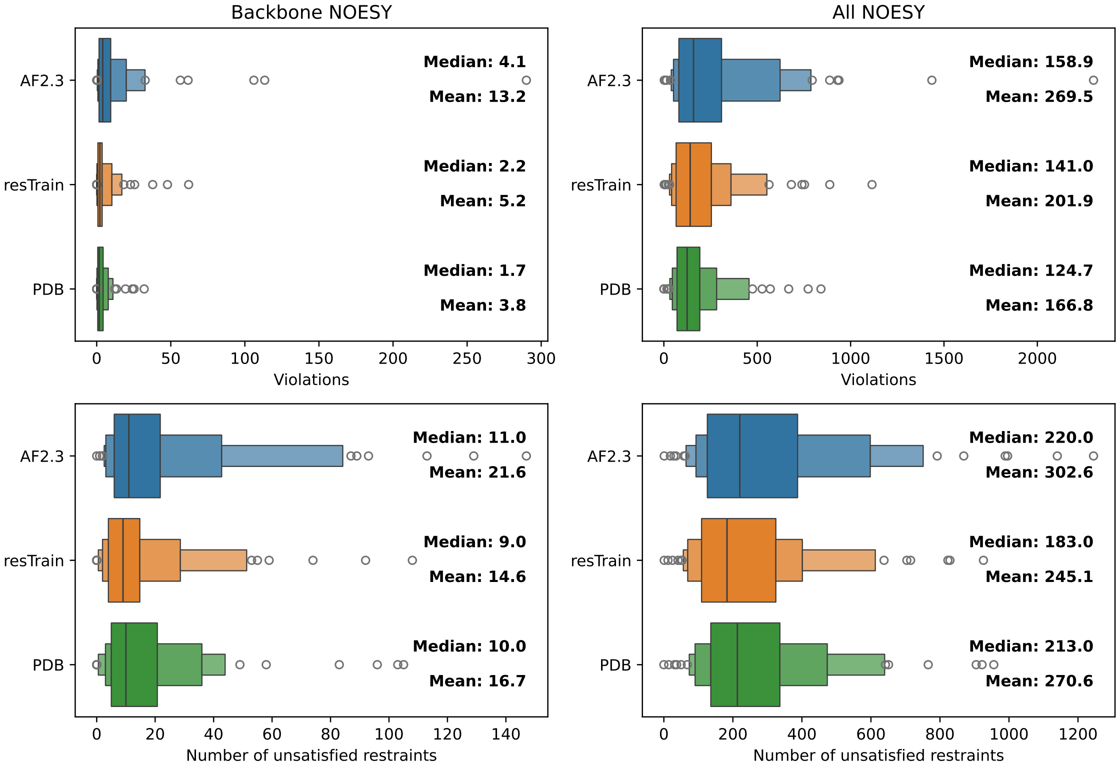


Figure 2: Boxen plots of NOESY restraint violations. Violations are calculated either against NOESY restraints involving backbone H and Q atoms (left column) or on all restraints (right column). Violations are summarised either as the aggregate (sum) of all errors (top row) or as the number of restraints that are not satisfied (bottom row). AF-resTrain performs better than AF2.3 across all metrics. AF-resTrain predictions violate fewer restraints on average than the corresponding best model deposited in the PDB, but the aggregate errors are larger.

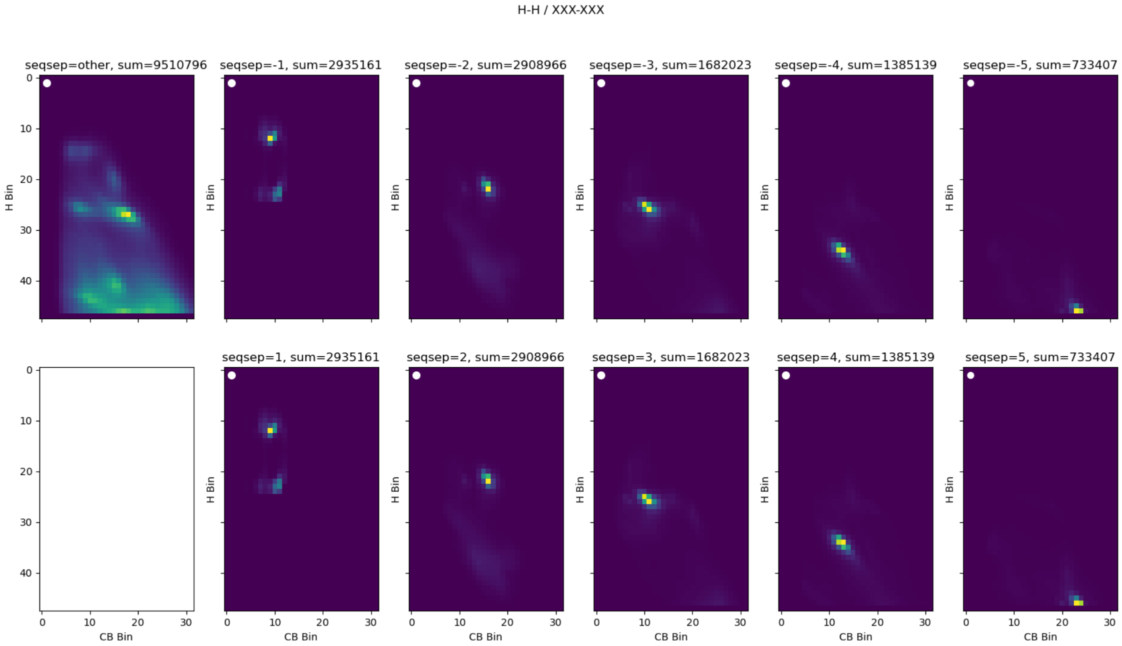


(a)

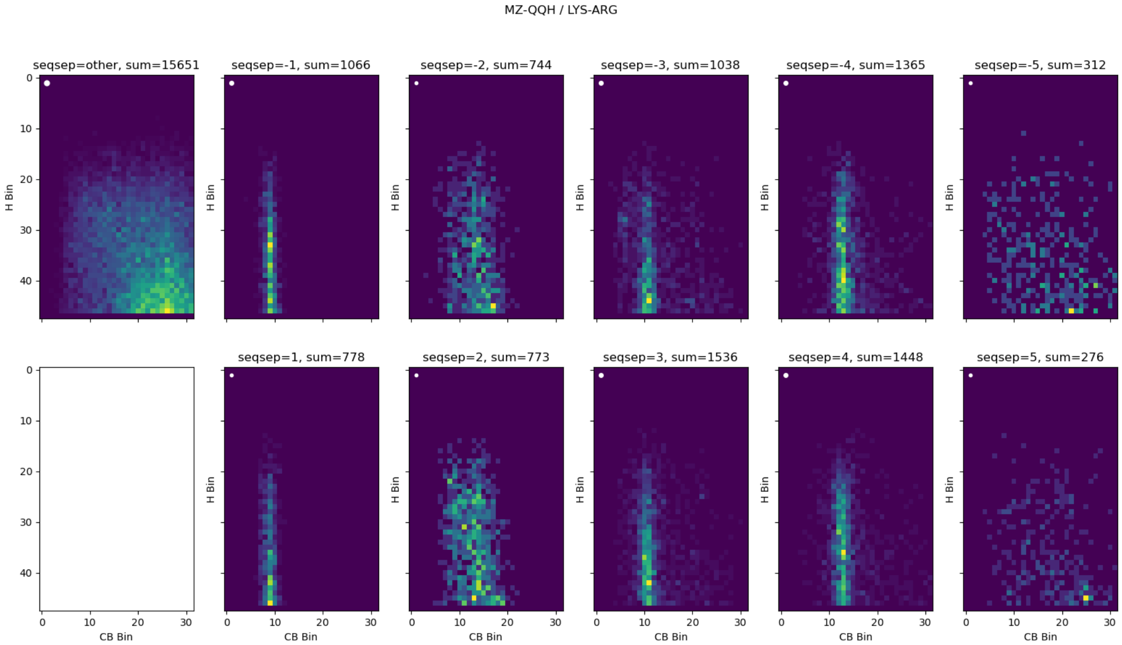


(b)

Figure 3: Example distributions of H-H distances for (a) ”H”-”H” and (b) ”QB”-”QQR” atoms and the corresponding CB-CB distance. Different panels show different sequence separation between the residues. Total number of counts per image is shown. (a) Preference for trans conformations can be seen for seqsep=$\pm 1$. (Prolines do not have an ”H” atom).

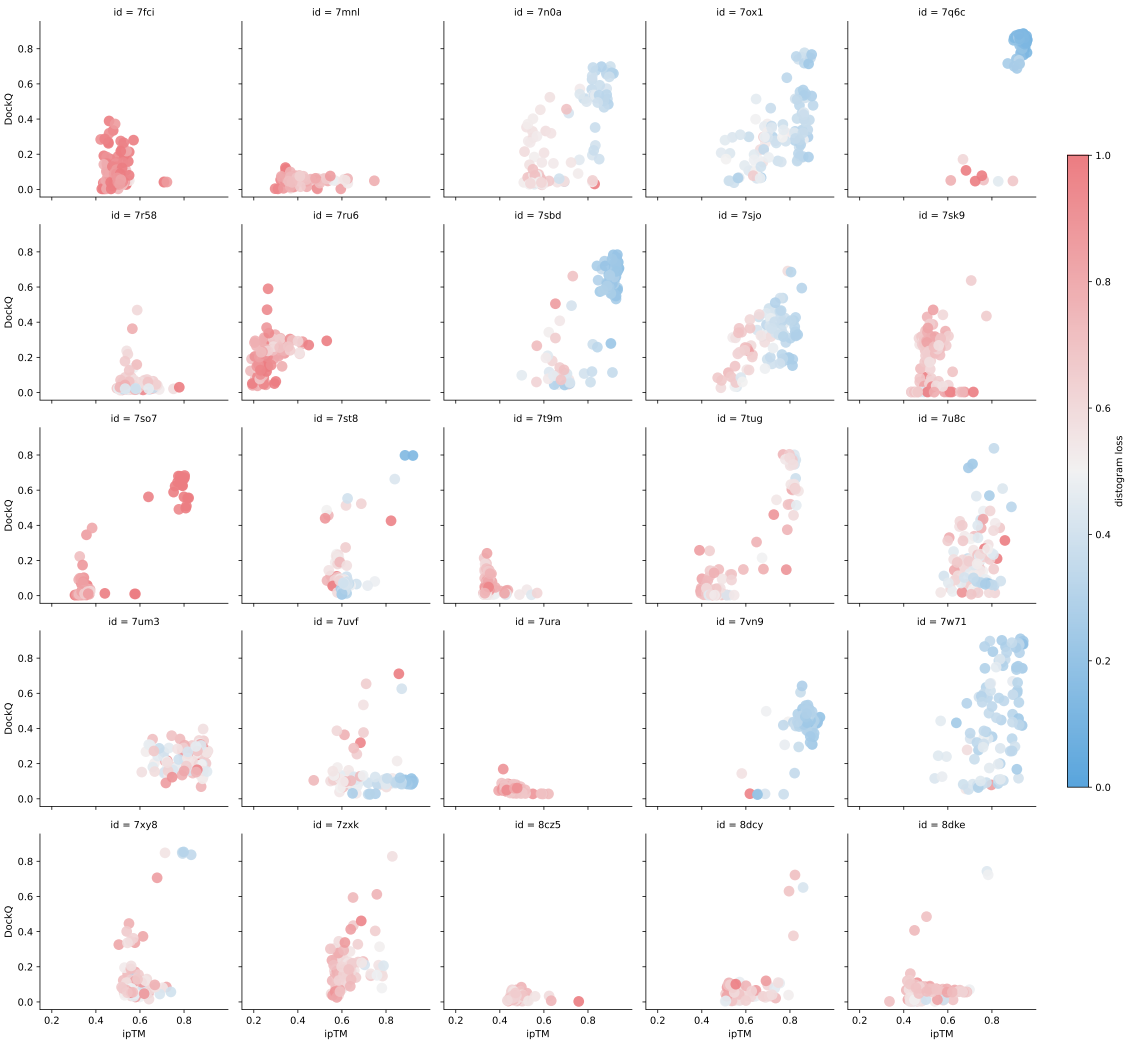


Figure 4: Scatter plots comparing predicted interface quality (ipTM) in AF2-resTrain with prediction quality measured by DockQ in the antigen-antibody benchmark. Points are colored by normalised distogram loss. For many of the targets, the predicted quality measure is able to correctly pick the higher-quality predictions.

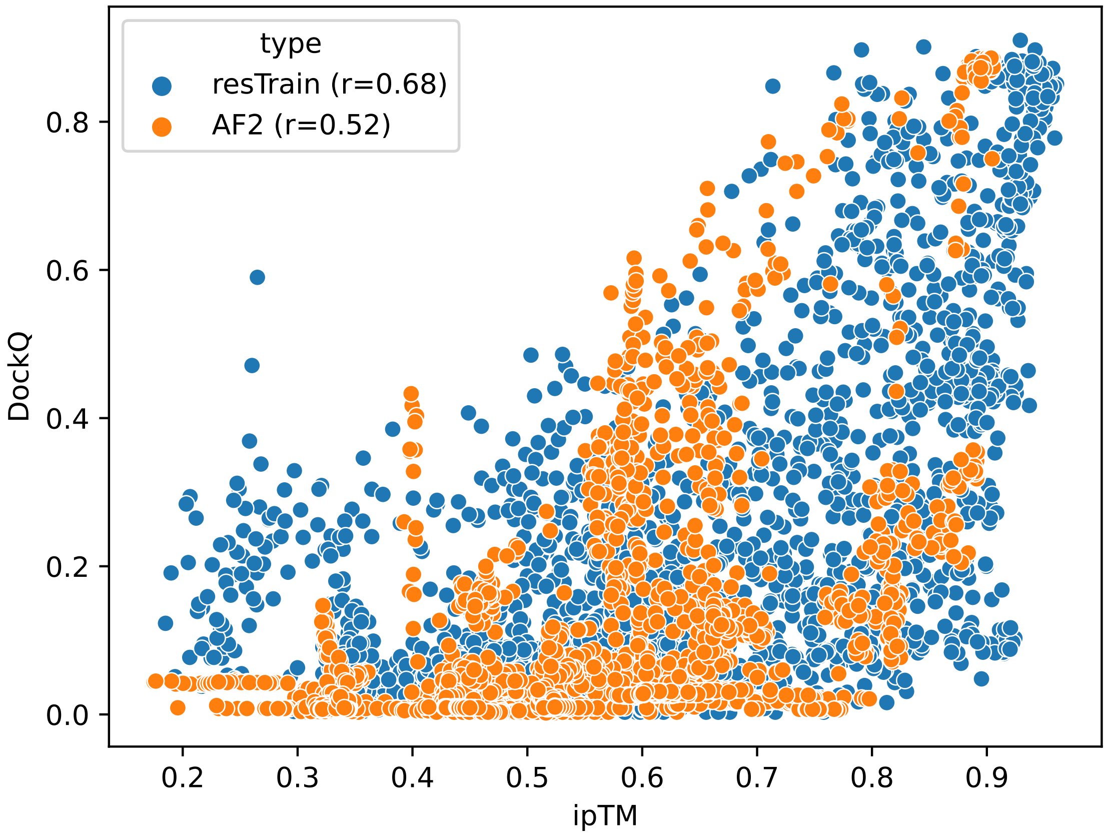


Figure 5: Scatter plots comparing correlation between ipTM and DockQ scores for AF2 and resTrain across all predictions of antigen-antibody benchmark.

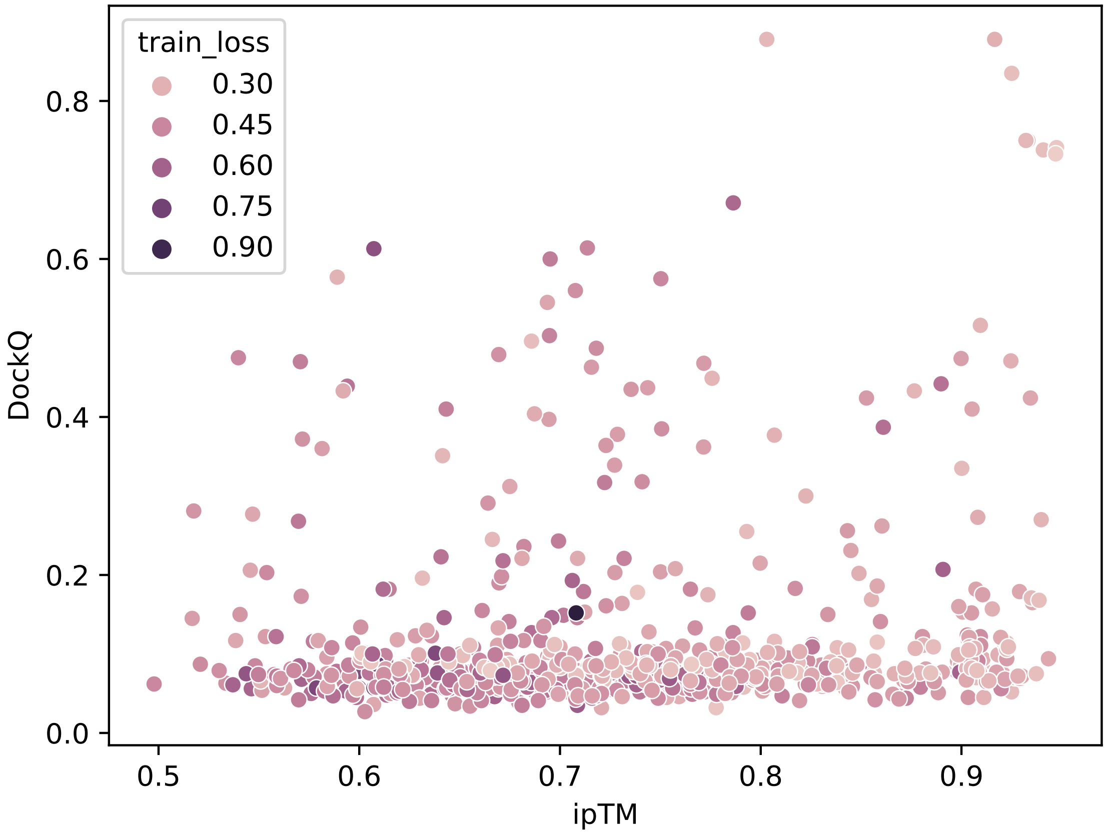


Figure 6: Scatter plots comparing predicted interface quality (ipTM) in AF2-resTrain with prediction quality measured by DockQ in the epitope scanning experiment (target 7W71). Multiple experiments are pooled together by forcing the CDR-H loop to be in contact with all receptor amino acids. The best overall model by ipTM is also one of the best-scoring models by DockQ.

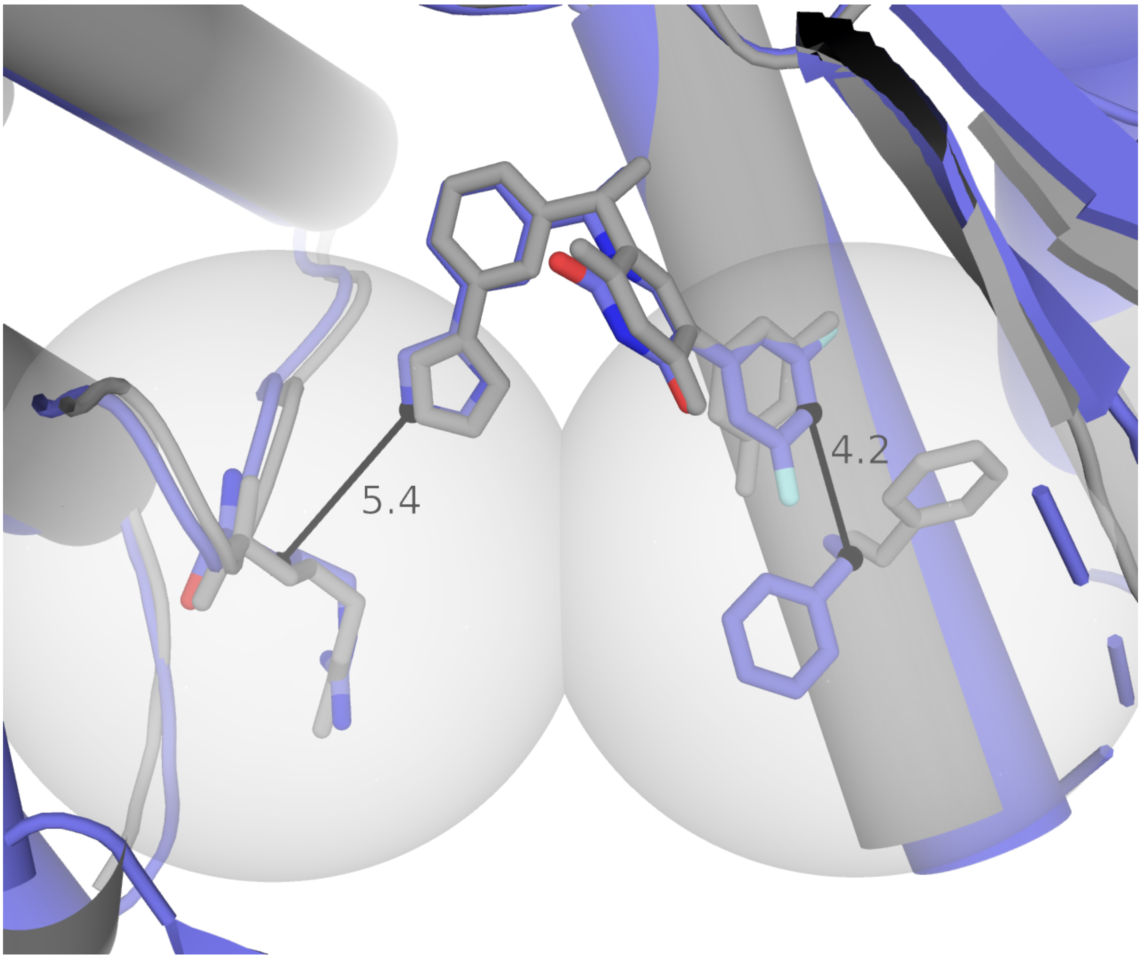


Figure 7: Example of aided protein-ligand docking with OF3-resTrain. We select two heavy atoms (N and C18, left and right, respectively) at either side of the ligand and their closest protein residues (represented by their CB atoms), then run OF3-resTrain by restraining the two pair distances to a maximum of 8Å (restraint radius around CB atoms depicted as spheres, black lines are labelled with distances in Å in best prediction). In the top ranked structure by ipTM, the ligand is correctly predicted (blue) when compared to the deposited structure (grey).

| Group | Pseudoatom | Hydrogens | Notes |
|---|---|---|---|
| Methyls | MB | HB1 HB2 HB3 | |
| | MG1 | HG11 HG12 HG13 | |
| | MG2 | HG21 HG22 HG23 | |
| | MD1 | HD11 HD12 HD13 | |
| | MD2 | HD21 HD22 HD23 | |
| | ME | HE1 HE2 HE3 | |
| | MZ | HZ1 HZ2 HZ3 | |
| Methylenes | QA | HA2 HA3 | |
| | QB | HB2 HB3 | |
| | QG | HG2 HG3 | |
| | QG1 | HG12 HG13 | ILE only |
| | QD | HD2 HD3 | |
| | QD2 | HD21 HD22 | ASN only |
| | QE | HE2 HE3 | |
| | QE2 | HE21 HE22 | GLN only |
| | QZ | HZ2 HZ3 | |
| | QH1 | HH11 HH12 | |
| | QH2 | HH21 HH22 | |
| Rings | QRD | HD1 HD2 | ARG only |
| | QRE | HE1 HE2 | ARG only |
| Composites | QQG | HG11 HG12 HG13 HG21 HG22 HG23 | |
| | QQD | HD11 HD12 HD13 HD21 HD22 HD23 | |
| | QQH | HH11 HH12 HH21 HH22 | |
| | QQR | HD1 HD2 HE1 HE2 | |

Table 1: Atom/pseudoatom mappings.

| Residue | (Pseudo)Atom | Mapped Class | Notes |
|---|---|---|---|
| Valine | MG1 | MGX_VAL | two chemically |
| | MG2 | MGX_VAL | equivalent groups |
| Leucine | MD1 | MDX_LEU | two chemically |
| | MD2 | MDX_LEU | equivalent groups |
| Asparagine | HD2 | HD2_ASP | unique atom type |
| Histidine | HD1 | HD1_HIS | unique atom type |
| | HD2 | HD2_HIS | unique atom type |
| | HE1 | HE1_HIS | unique atom type |
| | HE2 | HE2_HIS | unique atom type |
| Tryptophan | HD1 | HD1_TRP | unique atom type |
| | HE1 | HE1_TRP | unique atom type |
| | HE3 | HE3_TRP | unique atom type |
| | HZ2 | HZ2_TRP | unique atom type |
| | HZ3 | HZ3_TRP | unique atom type |
| | HH2 | HH2_TRP | unique atom type |
| Threonine | HG1 | HG1_THR | unique atom type |
| Glutamate | HE2 | HE2_GLU | unique atom type |
| Arginine | HE | HE_ARG | unique atom type |
| | QH1 | QHX_ARG | two chemically |
| | QH2 | QHX_ARG | equivalent groups |
| Phenylalanine | HZ | HZ_PHE | unique atom type |
| Tyrosine | HH | HH_TYR | unique atom type |

Table 2: Non-trivial atom/pseudoatom class mappings

| Variable | Binning |
|---|---|
| Hydrogen distance | Bin count: 48 bins<br>First bin lower bound: 0.75<br>Last bin lower bound: 8.0<br>Binwidth: approx 0.154Å |
| $C_\beta$ distance | Bin count: 64 bins<br>First bin lower bound: 2.3125<br>Last bin lower bound: 21.6875<br>Binwidth: approx 0.308Å |
| Sequence distance | -5, -4, -3, -2, -1, 1, 2, 3, 4, 5, other |

Table 3: H-H to $C_\beta$-$C_\beta$ distance conversion is achieved by aggregating and binning count data from the PDB. CB-CB atom distances are binned as in AlphaFold distograms, while the corresponding distances between H atoms are split into 48 bins in steps of approx. 0.154Å. Counts are also binned by sequence separation, calculated as the difference between the residue pair indexes on the protein sequence.
